# Female baboon adrenal zona fasciculata and zona reticularis regulatory and functional proteins decrease across the life course

**DOI:** 10.1101/2023.09.06.556552

**Authors:** Hillary F Huber, Cun Li, Dongbin Xie, Kenneth G Gerow, Thomas C Register, Carol A. Shively, Laura A. Cox, Peter W Nathanielsz

**Affiliations:** Southwest National Primate Research Center, Texas Biomedical Research Institute, San Antonio, TX, USA; Texas Pregnancy & Life-course Health Research Center, Animal Science, Univ of Wyoming, Laramie, WY, USA; Statistics, Univ of Wyoming, Laramie, WY, USA; Pathology- Comparative Medicine, Wake Forest School of Medicine, Winston-Salem, NC, USA; Center for Precision Medicine, Wake Forest School of Medicine, Winston-Salem, NC, USA

**Keywords:** Adrenocortical aging, Adrenocortical life-course function, nonhuman primate, cortisol, glucocorticoid, hypothalamo-pituitary-adrenal-axis

## Abstract

Debate exists on life-course adrenocortical zonal function trajectories. Rapid, phasic blood steroid concentration changes, such as circadian rhythms and acute stress responses, complicate quantification. To avoid pitfalls and account for life-stage changes in adrenocortical activity indices, we quantified zonae fasciculata (ZF) and reticularis (ZR) across the life-course, by immunohistochemistry of key regulatory and functional proteins.

In 28 female baboon adrenals (7.5-22.1 years), we quantified 12 key proteins involved in cell metabolism, division, proliferation, steroidogenesis (including steroid acute regulatory protein, StAR), oxidative stress, and glucocorticoid and mitochondrial function.

Life-course abundance of ten ZF proteins decreased with age. Cell cycle inhibitor and oxidative stress markers increased. Seven of the 12 proteins changed in the same direction for ZR and ZF. Importantly, ZF StAR decreased while ZR StAR was unchanged. Findings indicate ZF function decreased, and less markedly ZR function, with age. Causes and aging consequences of these changes remain to be determined.

## INTRODUCTION

Adrenocorticosteroids regulate multiple, key cellular functions across the life course including metabolic function [1] and related energy production and oxidative stress [1–3]. Both high circulating glucocorticoid concentrations that occur in Cushing’s syndrome and low concentrations that occur in Addison’s disease are accompanied by premature aging [1]. We have researched adrenocortical function in baboons, rhesus monkeys, sheep, and rats for various life course stages from fetal life to old age [3–6]. There is considerable debate on the trajectory of life-course changes in circulating adrenocorticosteroids. The lack of a robust picture of life-course adrenocortical activity is due in part to practical problems caused by rapid, often marked changes in blood adrenocortical steroid concentrations. These changes are a consequence of normal adrenocortical rhythms such as circadian and ultradian rhythms and, even more importantly, changes in responses to multiple different stress and disease challenges. Adrenocortical changes are more controversial during the aging stage of life than at any other time in the lifespan.

Obtaining repetitive blood samples over even short periods of time constitutes a stress to the hypothalamo-pituitary-adrenal-axis (HPAA), even when chronic indwelling catheters are in place [4,7]. As a result, very few normative studies provide data on changes of circulating concentrations of cortisol and dehydroepiandrosterone (DHEA) across significant portions of the life course in any mammalian species [6–9]. This lack of information is particularly true in primate species. In primates, cortisol is produced in the adrenal zona fasciculata (ZF) and DHEA in the adrenal zona reticularis (ZR) [3,8].

Unfortunately, most life-course studies on changes in blood steroid concentrations are categorical analyses based on blood values obtained at a few, sometimes only two, age timepoints [3,9–11]. In humans, nonhuman primates, and rodents, available reports show a variety of results with glucocorticoids increasing, falling, or showing no change with age [3]. Because of the many roles cortisol and DHEA play in physiological processes involved in aging and health span, it is important to determine a complete profile of life-course steroid concentrations. To enable translation to human biology it is particularly important that nonhuman primate data are available. Since studies are not permitted on great ape species, the baboon is the experimental nonhuman primate species with the closest physiology to humans [12]. Two recent, completely independent baboon studies of separate colonies maintained at different locations described a linear fall in baboon serum cortisol from ∼5-23 years (lifespan ∼21y) [9,11]. Slopes of the linear falls were remarkably similar: -24.7 and -23.7 ng. cortisol ml ^-1^.y^-1^.

We have studied several physiological systems in a unique cohort of baboons maintained in external group housing allowing free physical and social interaction at the Southwest National Primate Research Center (SNPRC) in San Antonio Texas for over 30 years [9,13–15]. Twenty-eight females from this cohort underwent necropsy at 7.5-22.1 years of age. We were unable to include males across the life course as primate centers do not maintain adequate numbers of males beyond the early years of life. The published lifespan for baboons in our colony is 21 years [16]. As a different, and potentially more integrated approach to evaluating life-course changes, we quantified 12 key regulatory and functional proteins in adrenocortical function by immunohistochemistry in the ZF and ZR of adrenals removed across the life course from these 28 female baboons. These 12 proteins play key regulatory or functional roles in the adrenal cortex across the life course. We propose that zonal content of these regulatory proteins in the ZF and ZR is a good surrogate for circulating adrenocorticosteroid levels and minimizes phasic changes in production caused by altered biology in different periods of life such as puberty, due to circadian rhythms and stress related rapid environmental changes in steroid concentrations. Importantly, since the adrenal cortex is a complex, heterogeneous organ containing multiple cell types, a histological approach enables evaluation of the spatial content of specific proteins in specific zones. In contrast, Western blot analysis undertaken on the adrenal or even adrenocortical homogenates inevitably includes cell types from zones in which analysis is not sought. To study overall cellular metabolism, we measured mechanistic target of rapamycin (mTOR). For cell division and proliferation, we measured cyclin-dependent kinase 4 (CDK4), cyclin-dependent kinase 1/2 (CDK1,2), protein Ki67 (Ki67), proliferating cell nuclear antigen (PCNA), and cyclin-dependent kinase inhibitor 1 (P21). For steroidogenesis, we measured steroidogenic acute regulatory protein (StAR). For glucocorticoid function, we measured glucocorticoid receptor (GR). For mitochondrial function, we measured citrate synthase (CS), cyclophilin D (CPY40), and ATP-synthase (ATP5A). Finally, to evaluate oxidative stress, we measured nitrotyrosine (NT).

We hypothesized that a significant decrease in key protein factors that stimulate activities central to serum cortisol and DHEA production in these two zones would provide evidence to support an age-related decreased adrenocortical function. Conversely an increase in proteins that inhibit steroid production also would indicate decreased function.

## METHODS

### Animal care and maintenance

Baboons (*Papio* sp., n=28 females ages 7.5-22.1 yrs; approximate human age equivalent 30-88 yrs) were housed in outdoor group cages at SNPRC, located at Texas Biomedical Research Institute (Texas Biomed) in San Antonio, TX. All animal procedures were approved by the Texas Biomed Institutional Animal Care and Use Committee (IACUC) and conducted in facilities approved by the Association for Assessment and Accreditation of Laboratory Animal Care (AAALAC) International. Diet was normal monkey laboratory chow (Purina 5LE0, containing 15% protein, 4% fat, 10% fiber, 0.22% glucose, 0.24% fructose, and metabolizable energy content of 2.98 kcal/g; Purina, St Louis, MO, USA) *ad libitum*, a complete life-cycle diet for Old World primates. Water was continuously available. SNPRC veterinarians conduct full physical exams twice annually and whenever else required. All animals were given a full veterinary examination prior to recruitment to the study and were in good health at time of study.

### Necropsy to obtain adrenals and other tissues

We studied adrenals from 28 female baboons across the life course at 7.5-22.1 years. After an overnight fast, baboons were sedated with ketamine (10 mg/kg), and then anesthetized with 1-2% isoflurane. While under general anesthesia baboons were euthanized by exsanguination as approved by the American Veterinary Medical Association. Following cardiac asystole, all tissues were collected according to a standardized protocol within <1 hr from time of death, between 8:00-10:00 AM to minimize potential variation from circadian rhythms. Adrenal weights, body weights, and body morphometrics are shown in **Table 1**. We have shown that euthanasia by intravenous agents can alter tissue structure [17].

**Table 1.** Baboon ages, body weights, body morphometrics, and adrenal weights. The few missing data were due to the absence of technical staff.

| Animal | Age (yrs) | Body weight (kg) | Body length (cm) | Body mass index | Chest circ (cm) | Waist circ (cm) | Hips circ (cm) | Left adrenal weight (g) | Right adrenal weight (g) |
| --- | --- | --- | --- | --- | --- | --- | --- | --- | --- |
| 1 | 7.5 |  |  |  |  |  |  |  |  |
| 2 | 8.0 | 14.3 |  |  |  |  |  |  |  |
| 3 | 8.7 | 16.1 | 102.5 | 15.4 | 52.2 | 46.2 | 42.5 | 0.8 | 0.7 |
| 4 | 8.9 | 19.0 | 104.0 | 17.5 | 53.2 | 47.8 | 44.3 | 0.9 | 1.1 |
| 5 | 9.4 | 21.9 |  |  |  |  |  |  |  |
| 6 | 10.2 | 16.5 |  |  |  |  |  |  |  |
| 7 | 10.4 | 18.6 | 100.0 | 18.6 | 56.9 | 64.1 | 49.0 | 1.3 | 1.1 |
| 8 | 11.9 | 17.2 | 111.0 | 14.0 | 50.5 | 43.0 | 38.0 | 1.4 | 1.1 |
| 9 | 12.0 | 18.5 | 108.0 | 15.9 | 56.0 | 43.0 | 44.5 | 1.3 | 1.2 |
| 10 | 12.9 | 17.5 | 107.0 | 15.3 | 50.6 | 45.0 | 43.5 | 1.3 | 1.0 |
| 11 | 13.2 | 17.5 | 103.0 | 16.5 | 55.2 | 51.3 | 46.0 | 1.0 | 0.9 |
| 12 | 13.6 | 17.5 | 104.5 | 16.0 | 51.7 | 41.7 | 37.3 | 0.8 | 0.6 |
| 13 | 13.8 | 17.8 | 107.3 | 15.5 | 57.8 | 50.4 | 26.7 | 1.3 | 1.9 |
| 14 | 14.8 | 18.4 | 112.0 | 14.7 | 55.0 | 43.0 | 51.0 | 1.0 | 0.8 |
| 15 | 15.1 | 20.2 | 105.5 | 18.1 | 59.5 | 56.5 | 45.2 | 0.6 | 0.7 |
| 16 | 15.7 | 17.0 | 106.5 | 15.0 | 55.0 | 48.0 | 47.0 | 0.9 | 0.6 |
| 17 | 17.9 | 20.3 | 108.7 | 17.2 | 62.8 | 64.6 | 54.9 | 0.9 | 0.8 |
| 18 | 17.9 |  |  |  |  |  |  |  |  |
| 19 | 18.3 | 30.9 | 110.5 | 25.3 | 81.2 | 85.6 | 66.0 | 1.4 | 1.3 |
| 20 | 18.4 | 16.6 | 105.0 | 15.1 | 57.6 | 49.6 | 48.7 | 0.9 | 0.8 |
| 21 | 18.5 | 16.2 | 109.5 | 13.5 | 53.4 | 41.0 | 39.0 | 1.4 | 1.1 |
| 22 | 18.6 | 25.0 | 103.5 | 23.3 | 73.5 | 77.2 | 63.0 | 1.3 | 1.1 |
| 23 | 18.9 | 23.8 | 105.5 | 21.4 | 67.2 | 68.2 | 52.0 | 0.9 | 0.7 |
| 24 | 19.2 | 17.6 | 106.5 | 15.5 | 50.8 | 54.5 | 42.7 | 1.1 | 0.8 |
| 25 | 19.8 | 16.2 | 99.0 | 16.5 | 61.4 | 49.6 | 49.2 | 1.1 | 0.9 |
| 26 | 20.8 | 15.1 | 97.5 | 15.9 | 53.0 | 51.0 | 40.0 | 0.9 | 0.8 |
| 27 | 21.3 | 13.7 | 95.0 | 15.2 | 48.7 | 55.6 | 37.0 | 0.8 | 0.6 |
| 28 | 22.1 | 17.6 | 93.3 | 20.2 | 55.8 | 58.4 | 46.0 |  | 0.8 |

### Histology and quantitative immunohistochemistry

We studied 12 regulatory and functional proteins involved in steroidogenesis in both the ZF and ZR (**Table 2**). Adrenal glands were fixed for 48 hours with 4% paraformaldehyde solution, dehydrated, and blocked in paraffin. Three 5 μm sections taken at 150 m intervals from the region opposite the vascular pole of the adrenal were immunostained for CS, StAR, mTOR, Cdk4, CDK1,2, CPY40, Ki67, NT, GR, P21, PCNA, and ATP5A. Sources and final concentrations of the primary antibodies for the proteins together with validation procedures – western blot and negative control – are presented in Table 2. Immunohistochemical methods have been described in detail [18]. The final optimal concentration of the primary antibodies for the proteins was determined using our published methods [19]. For zone demarcation, since we use paraffin sections in which we cannot see lipid droplets well, we demarcated based on the arrangement of cells. The ZF appears as parallel cords or columns of larger cells, while the ZR shows irregular cords or clusters of smaller cells. This distinct cellular arrangement helps differentiate between the two regions.

**Table 2.** Antibody antigens, final dilutions, and sources used for immunohistochemical staining of baboon adrenal tissue.

| **Antibody** | **Company** | **Cat #** | **Concentration** | **Negative Control** | **Data Sheet** |
| --- | --- | --- | --- | --- | --- |
| Citrate synthase (CS) | Santa Cruz | sc-390693 | 1:150 | mouse IgG cell | sc-390693 (scbt.com) |
| Steroidogenic acute regulatory protein (StAR) | Santa Cruz | sc-166821 | 1:2000 | mouse IgG cell | sc-166821 (scbt.com) |
| Mechanistic target of rapamycin (mTOR) | Santa Cruz | sc-517464 | 1:50 | mouse IgG cell | SCBT |
| Cyclin-dependent kinase 4 (Cdk4) | Santa Cruz | sc-166373 | 1:100 | mouse IgG cell | sc-166373 (scbt.com) |
| Cyclin-dependent kinase 1/2 (Cdk1,2) | Santa Cruz | sc-53219 | 1:50 | mouse IgG cell | sc-53219 (scbt.com) |
| Cyclophilin D (CPY40) | abcam | ab181983 | 1:500 | Normal rabbit serum | datasheet_181983 (2).pdf |
| Protein Ki67 (Ki67) | Novocastra | NCL-ki67-mm1 | 1:50 | mouse IgG cell | Novocastra Ki67 Antibody for IHC Staining (leicabiosystems.com) |
| Nitrotyrosine (NT) | Santa Cruz | sc-32757 | 1:500 | mouse IgG cell | sc-32757 (scbt.com) |
| Glucocorticoid receptor (GR) | Santa Cruz | sc-393232 | 1:50 | mouse IgG cell | sc-393232.qxp (scbt.com) |
| Cyclin-dependent kinase inhibitor 1 (P21) | Santa Cruz | sc-817 | 1:150 | mouse IgG cell | sc-817 (scbt.com) |
| Proliferating cell nuclear antigen (PCNA) | Santa Cruz | sc-56 | 1:1000 | mouse IgG cell | sc-56 (scbt.com) |
| ATP-synthase (ATP5A) | Santa Cruz | sc-136178 | 1:200 | mouse IgG cell | sc-136178 (scbt.com) |

Images were obtained with Nikon Ri2 Color Camera and imaging with Nikon NIS Elements D Software. Six pictures (2650 x 1920 pixels) were taken at the 2, 4, 6, 8, 10, and 12 O’clock section positions and analyzed with NIH Image J software for fraction (area immunostained / area of the field of interest x 100%).

### Statistical analysis of life course changes in protein abundance

To determine effects of age on all proteins measured, data for each protein were log transformed to exploit the essential exponential decay (or increase) that was visible in most cases and regressed against age, using a simple linear regression model. In turn, back-transforming the slope coefficients yields rates of change, e.g. 10% increase per year. Regarding percent changes, for example, a 10% increase from one year to the next starting at 20% area stained becomes 22%. Significant differences between rates of change for each protein in ZF and ZR were verified by a two-sample t-test. Significance was set at P < 0.05.

## RESULTS

Protein immunoreactivity changed with age in both the ZF and ZR. Examples of immunostaining differences between a young adult 9-year-old and elderly 21-year-old are shown in Figure 1 for ZF and Figure 2 for ZR. In ZF, 10 of the 12 proteins deceased with age (Figure 3). The exceptions were NT and P21, which both increased with age; these two proteins also increased with age in ZR. In ZR, CDK1,2, Ki67, GR, PCNA, and ATP5A decreased with age, while the remaining 5 proteins did not change with age (Figure 3). Rates of increase and decrease in proteins with each year of aging are shown in Table 3.

**Table 3.** Yearly percentage change in staining (% area of tissue stained) for each of the 12 proteins in the ZF and ZR. A negative number indicates a fall in protein staining with age; a positive number indicates an increase. The p values for the rates of fall are given with each protein in **Figure 3**. The differences between the two rates of fall are also shown in **Figure 3**. *p<0.05; #p=0.08

|  |  | Zona Fasciculata | Zona Reticularis |
| --- | --- | --- | --- |
| Markers |  | rate % per year | rate % per year |
| Cell division and Proliferation | PCNA | - 8.4* | - 14.7* |
|  | Cdk4 | - 17.8* | - 4.6# |
|  | Cdk1,2 | - 17.5* | - 16.2* |
|  | Ki67 | - 16.8* | - 13.6* |
|  | P21 | + 16.3* | + 16.3* |
| Steroidogenesis | StAR | - 21.2* | 0.0 |
| Glucocorticoid Function | GR | - 15.2* | - 13.5* |
| Oxidative Stress | NT | + 19.9* | + 16.1* |
| Mitochondrial function | CS | - 22.9* | - 3.1 |
|  | CPY40 | -17.0* | - 1.8 |
|  | ATP5A | - 8.4* | - 12.5* |
| Overall Cellular Metabolism | mTOR | - 19.0* | - 3.6 |

**Figure 1.**
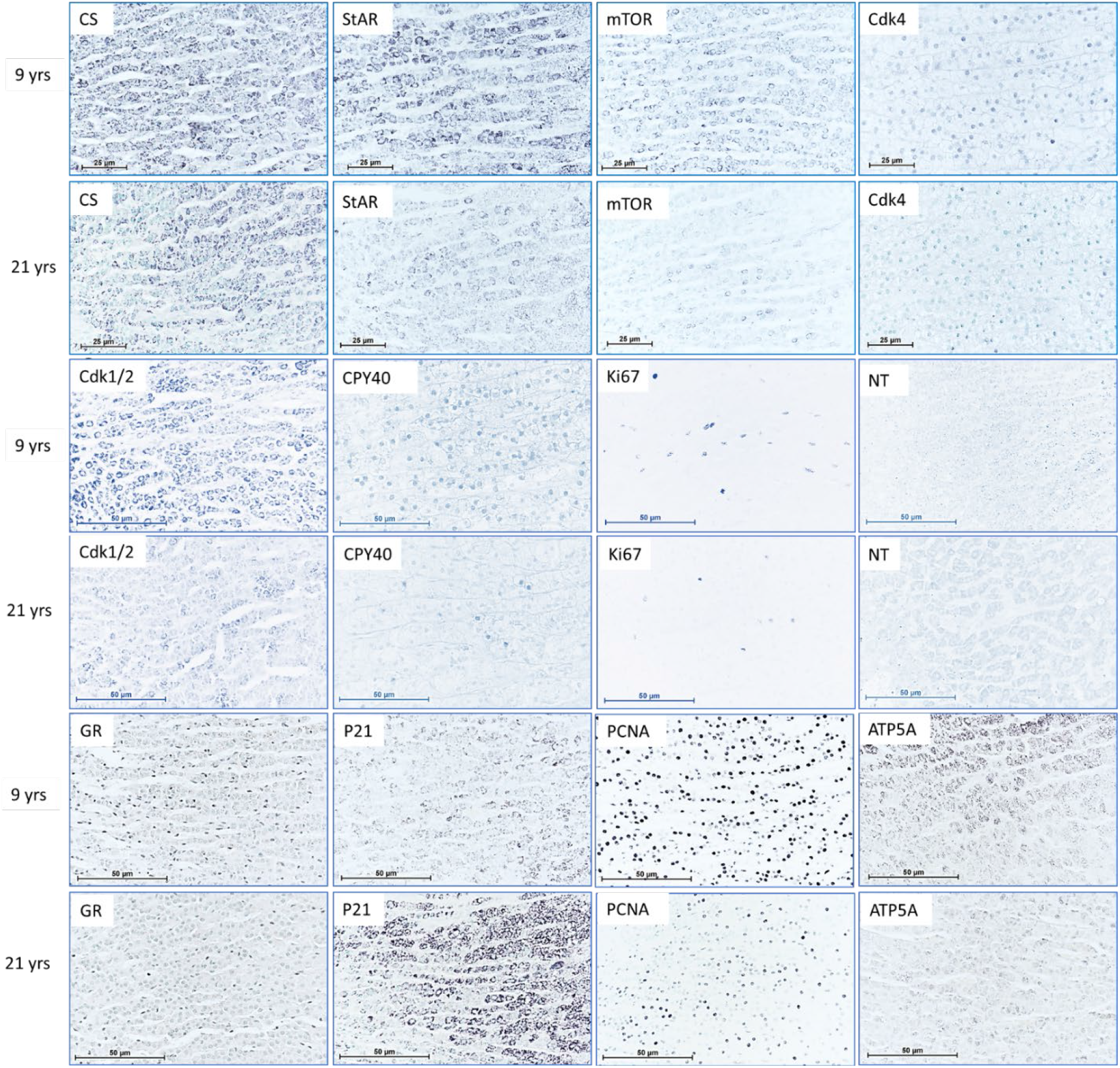
Immunostaining in the ZF for all 12 proteins at the extremes of the age range -a young age of 8.9 years and an old age of 21.3 years

**Figure 2.**
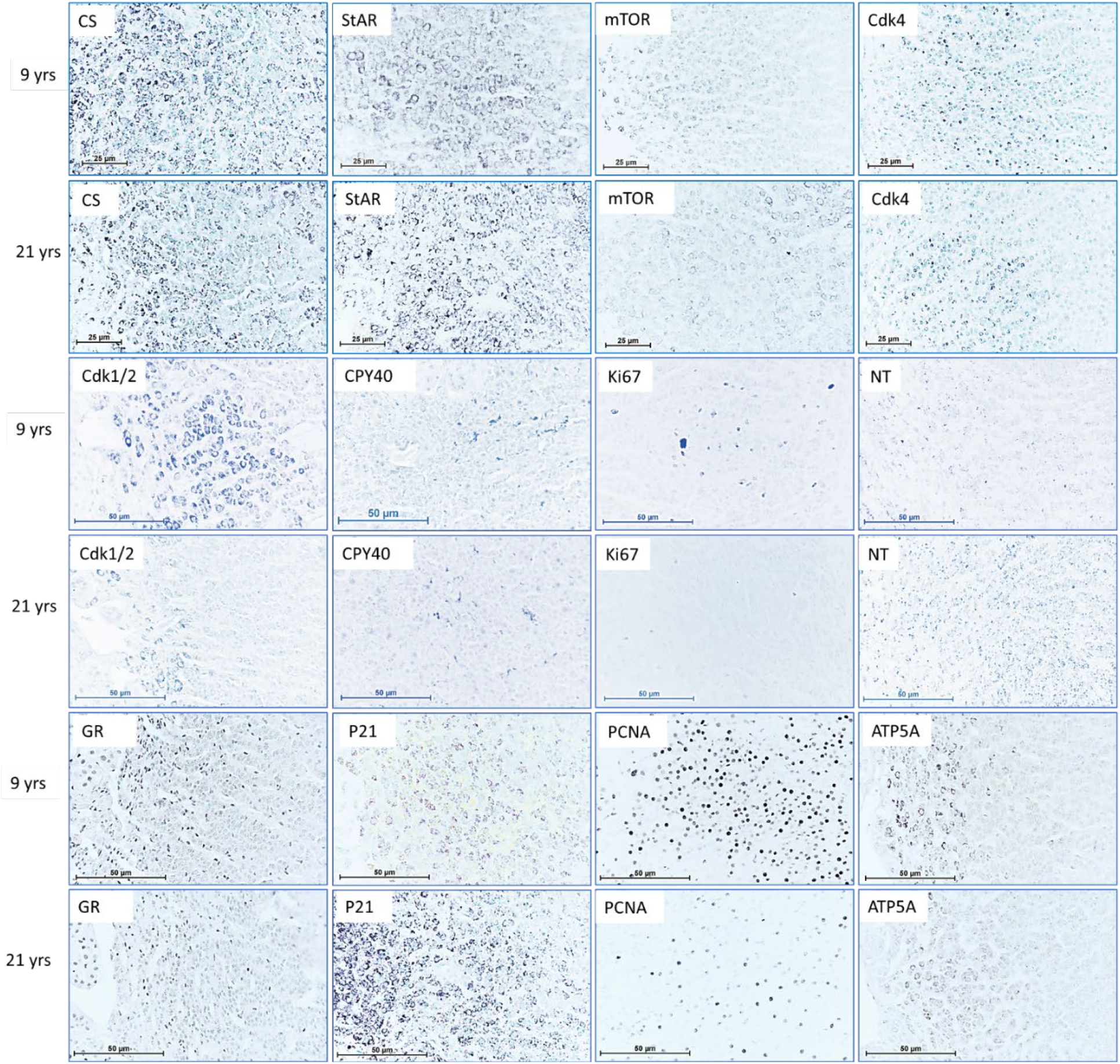
Immunostaining in the ZR for all twelve proteins at the extremes of the age range -a young age of 8.9 years and an old age of 21.3 years

**Figure 3.**
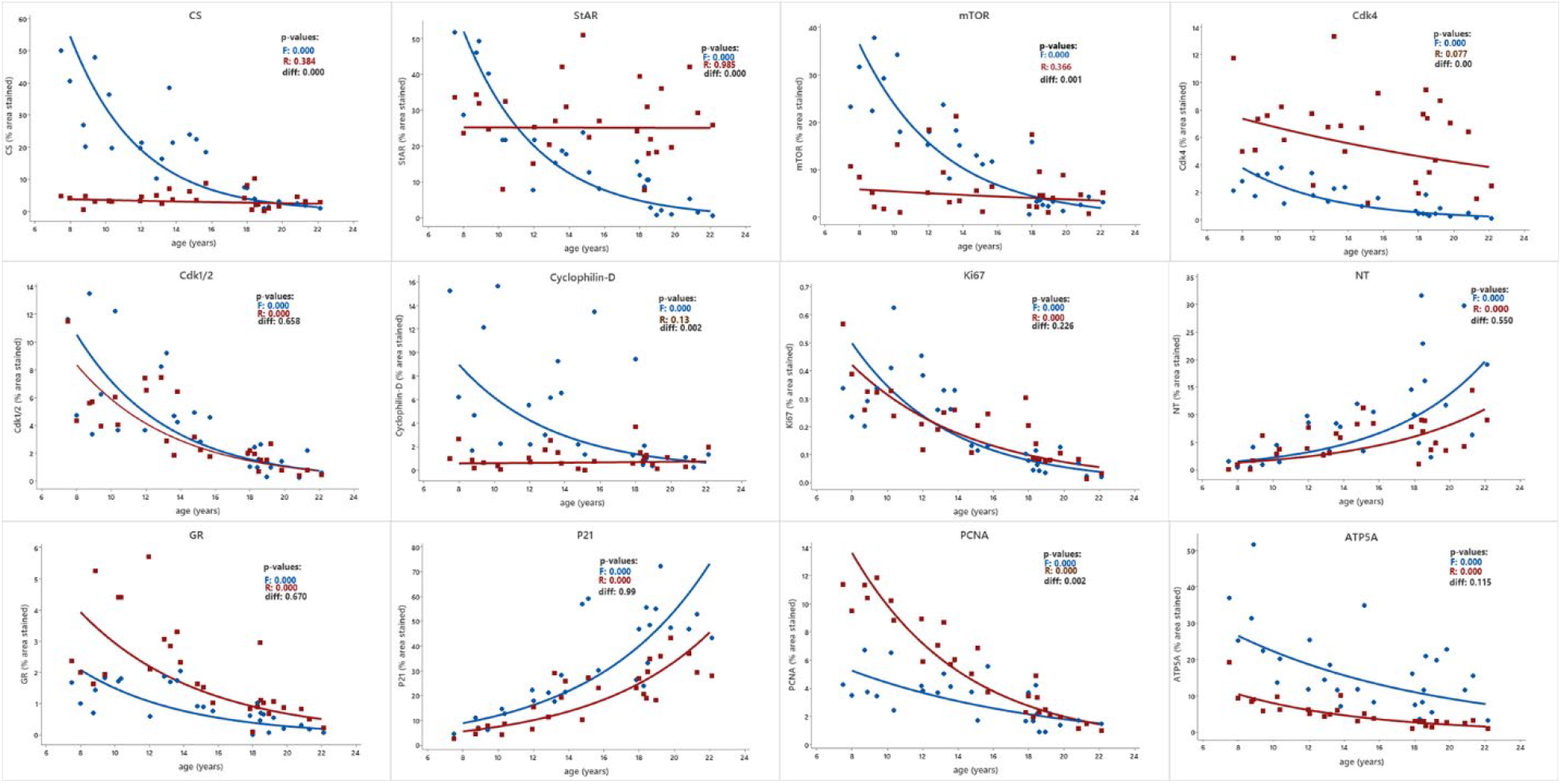
Percentage annual change in area of tissue stained for all 12 proteins in the adrenal ZF (blue) and ZR (red) of 28 female baboons

## DISCUSSION

### The importance of studies that address life-course ZF and ZR function

Multiple key metabolic adrenocorticosteroid functions are essential for normal cellular function throughout life. Both the extent and nature of adrenocorticosteroid function likely change at different rates at different stages of the life course. It is, therefore, an important first step in understanding the role of adrenocorticosteroids across the life-course, including frailty and aging, to establish the precise timing of life-course changes in ZF and ZR cellular activity in relation to normal aging. In rats we have shown a sexually dimorphic rise in corticosterone in the first half of life followed by a progressive fall [8]. We identified some limitations in the available life-course data in rodents and other species. First, in most adrenocorticosteroid life-course studies data are only presented at a few categorical ages, generally two or three [8]. It is essential that robust life-course datasets contain values from as many ages across the lifespan as practically possible to permit detailed regression rather than just two-point analysis. Second, well-accepted markers of aging and frailty, such as grip strength, are changing as early as one-third of the way through the lifespan [20]. Therefore, to understand the antecedents of aging adrenocortical function, data are needed well before any of the known hallmarks of aging emerge.

The baboon lifespan has been reported as 21 yrs in a female-only cohort at SNPRC [16]. In an ongoing, unpublished multi-institutional Nonhuman Primate Lifespan Project, we have preliminary data showing a maximum lifespan of 30.6 yrs, with 50% of baboons dying of natural causes or euthanized for health/clinical reasons by 11.5 yrs in females and 10.6 yrs males, and 95% dying by 21 yrs in both sexes (n=698 females, 393 males). Female baboons are considered fully grown at age 7 yrs [21]. Therefore, our sample from 7.5-22.1 years covers the entire baboon adult lifespan after maximum growth.

In a recent paper reporting life-course changes in circulating rodent corticosterone, we detailed the conflicting literature on circulating rodent glucocorticoid concentrations [3]. Prior to that study, as mentioned above, the primate center at Oklahoma University had shown a life-course fall in early morning fasting cortisol using data beginning around the time of puberty in male and female baboons, with a slope of -23.7 ng. cortisol. ml ^-1^. y^-1^ [11]. We have conducted the same measurements in female baboons over the same portion of the lifespan. The slope of the fall in our independent study was -24.7 ng. cortisol ml ^-1^.y^-1^.[9] The similarity in the independently obtained data are remarkable. One human study measured cortisol in four hourly blood samples obtained over one day at 19-25, 42-65, and 66-89 years in 44 men and 27 women. The results showed a negative correlation with age in plasma cortisol area under curve. The acrophase of the 24-hour cycle for cortisol was also negatively correlated with age [22]. The authors suggest that their results could be explained by a weakened response of the 24-hour HPAA system, which would be compatible with decreased ZF secretory capability. According to one major authority in aging research, most studies indicate that plasma total cortisol concentrations do not increase with advancing age in humans [23].

One of the major limitations in these studies is the lack of dynamic data to determine susceptibility to stress induced changes of blood cortisol concentrations at various physiological stages of life such as puberty, menopause, and aging phasic changes in the internal and external environments. The steroid blood concentrations are also affected by circadian and other rhythms. Therefore, as mentioned above, we propose that measuring ZF and ZR zonal content of key regulatory and functional proteins smooths out recent phasic concentration changes at each specific age under study.

### ZF changes across the life course

We found compelling evidence for decreased cell growth with age in the ZF. Of the five proteins associated with cell division and proliferation, the two known to have a role in increasing cell division, CDK4 and CDK1,2, showed a fall in immunoreactive protein staining intensity. CDK4 is a member of a family of cyclins that control cell division and proliferation that are fundamental cellular processes mediating progression through the G1 phase as the cell begins the process of DNA synthesis [24]. CDK1,2 is essential for cell transition from stage G2 to M. PCNA and Ki67 increase in dividing cells. Across the life course PCNA protein is synthesized in the early phases of the cell cycle (G1 and S) and plays an important role in cell cycle progression, DNA repair, and replication [25]. PCNA is commonly used as a marker of cell proliferation. Ki67 antigen protein is confined to the nucleus, is present at all stages of the cell cycle except G0, is associated with regulation of the cell cycle, and is absent in quiescent, non-dividing cells. P21 was the sole cyclin-related protein we measured whose abundance increased in the ZF with age. High P21 levels mediate G1 arrest via CDK inhibition [26], thereby regulating the balance between cell proliferation and quiescence in favor of quiescence. It is of interest that CDK4 and CDK1,2 both fall at approximately 17% per year and Ki67 falls and P21 increases at approximately 17% per year. Taken together these changes provide compelling evidence of decreased cell division and proliferation in the aging ZF.

StAR is present within mitochondria in all tissues that produce steroids [27–29]. StAR plays an essential role in steroid synthesis by regulating cholesterol transport from the outer to the inner mitochondrial membrane, making it available to the enzymes that convert cholesterol to the different adrenal steroids. StAR is thus the first step in the steroid synthesis pathway and is considered by many to be the rate-limiting step in steroidogenesis although this opinion is not held universally [28]. The marked fall in StAR with aging in the ZF in our study would lead to decreased synthesis of cortisol with aging. Of the proteins we measured, only CS (see below) falls at a faster rate than StAR with age. This observation adds strong support for the view that steroidogenesis falls with age.

To our knowledge very little information exists relevant to potential paracrine or autocrine feedback of glucocorticoids within the ZF. Entering the terms glucocorticoid receptor and zona fasciculata in PubMed produced only one report [29]. This paper indicated that it was the first report to demonstrate localization and distribution of GR throughout the rat adrenal cortex and reported that endogenous nitric oxide (NO) may modulate adrenocortical expression of this steroid receptor. If NO is a regulatory factor, that would support previous studies that show the splanchnic nerve influences adrenocortical steroid production [30].

Since steroids are mitochondrial products, we evaluated changes in NT, a marker of cell protein damage and nitric oxide production as a result of dysregulation of mitochondrial function. NT is produced by nitration of tyrosine. Increased NT is seen in many aging tissues [31]. Our data show a nearly 20% per year increase in ZF NT. CS is a mitochondrial matrix protein that is the initial and rate-limiting enzyme in the tricarboxylic acid cycle. CS actions are essential for mitochondrial respiration and are involved in glucose to lipid conversion. CS production is controlled by nuclear DNA. Following synthesis in the cytoplasm the enzyme is transported into the mitochondrion and is responsible for the formation of citrate in the citric acid cycle. CS is often used as a marker of mitochondrial number as its concentration increases with cellular activity. For example, the amount of the enzyme increases with muscular activity [32]. In support of the approach we have adopted, it is important to note that StAR and CS regulate steroidogenesis through different mitochondrial mechanisms and are the proteins whose staining falls the most with age [33]. CPY40 is located in the mitochondrial matrix and its staining also decreases with age. CPY40’s function has not been clearly defined and it may well have opposing functions at different times and concentrations. ATP5-A is a mitochondrial ATP-ase. Its decrease across the life course would lead to decreased energy production and accelerated aging.

mTOR is present in both mTOR complex 1 and mTOR complex 2. These two related proteins have multiple anabolic functions such as promoting cell growth and proliferation as well as protein synthesis. Thus, the pronounced decrease in mTOR protein in the ZF will lead to decreased ZF function.

In summary the changes observed in all 12 ZF proteins would lead to decreased cell growth, proliferation, protein synthesis, mitochondrial biogenesis, and metabolism with age. The most remarkable general feature of the changes in the ZF is the fall in overall staining in 10 of the 12 proteins. In two proteins, P21 and NT, the life course data show a rise in the staining. Except for P21, which is an inhibitor, these marked changes in proteins central to cell function and steroidogenesis indicate a pronounced fall in ZF mechanisms responsible for basal cortisol production.

### Comparison of protein life course changes in the aging ZF and ZR

We sought to determine whether there was evidence that the ZR and ZF age at different rates. While all 12 proteins studied showed significant changes in protein abundance across the life course in the ZF, only seven of the proteins showed a significant change in the ZR, and the direction was the same in the ZR and ZF. For six of the ZR proteins studied there was a significant decrease in protein abundance across the life course and the direction was the same in the ZR and ZF. In five of these the rate of decrease was greater in the ZF than ZR: CDK4, STaR, CS, CPY40, and mTOR. Abundance of two of the proteins, NT and P21, rose similarly in both ZR and ZF. The change in three of the proteins was significantly different in ZR from ZF. The most important of these was StAR, which remained unchanged across the life course in the ZR compared with a fall of -21.2% per year in the ZF. PCNA showed a greater positive slope in ZR than ZF, indicating a greater degree of preservation of nuclear number with age in the ZR than ZF. Together these results suggest a greater rate of aging in ZF than ZR.

Decreased mitochondrial function seems to play a larger role in ZF aging than ZR. StAR and citrate synthase regulate steroidogenesis through different mitochondrial mechanisms. Thus, it is of interest that their rates of fall in the ZF across the life course are very similar, while there was no significant fall in either protein in the ZR.

Obtaining life-course data in any species is demanding. Many practical problems account for the sparsity of life-course data available in the literature. Even in rats, subjects must be maintained for approaching three years, requiring a commitment of space, technician time, and adequate support funds. As mentioned above, we have published a study in which rat plasma corticosterone, the major glucocorticoid produced by the adrenal in rats, was measured at five points across the life course (postnatal days 21, 110, 450, 650, and 850). Lifespan in our colony is 850 days. We identified a significant fall in corticosterone between postnatal day 450 and 650 in both male and female rats [8]. This specific practical problem of frequent sampling across the life course is even more pronounced in nonhuman primates.

### Potential mechanisms responsible for the fall in ZF function across the life course

Adrenal steroid production by dispersed adrenal cells *in vitro* from 2, 5, 12, and 18-month-old rats show decreased age-related corticosterone production and adrenocorticotropic hormone (ACTH) responsiveness [34]. Ability to synthesize cholesterol for steroidogenesis by fresh male rat adrenal homogenates decreases with age. HMG-CoA activity was lowest at 12 and 18 months of age. Interestingly, this decrease is not associated with decreased activity of steroidogenic enzymes [17]. Corticosterone production by isolated male rat adrenal cells in response to ACTH and cyclic adenosine monophosphate (cAMP) has been studied from 6-24 months. Aged adrenocortical cells lose most of their ability to produce corticosterone in response to both ACTH and cAMP [35]. In this paper, the authors state, “Analysis of the data suggests that from 6 to 12 months, an intracellular steroidogenic lesion develops; in addition, there may be a loss in ACTH receptors on the plasma membrane. After 12 months these defects increase and are accompanied by a decrease in receptor sensitivity to ACTH.” A 6-12-month-old rat is equivalent to a 5-10-year-old baboon and an 15-30-year-old human. These data strongly support our findings of a fall in circulating corticosterone that begins around young adulthood [18].

One study reports greater protection against oxidative stress in 2-5-month-old rats than at 12-27 months. These data would fit with the age-related increase in NT in both the ZF and ZR [36]. While low oxidative stress levels and **r**edox signaling can be linked to normal cellular processes and are observed in normative aging, increased redox signalling is associated with age-related diseases [37]. Thus, our observation of a similar age-related increase in the marker of nitrosative stress in both the ZF and ZR would lead to increased cellular aging.

In summary, there are several strengths of this study of adrenocortical cellular markers of function across the life course. First, while many studies report data at limited categorical ages, our study uses regression analysis of data collected over the majority of the lifespan after puberty. We chose not to obtain measurements before puberty as there are so many rapid changes occurring in steroid function and general metabolism in that early developmental stage of life. Second, we have studied this cohort of baboons and their parents since 1985. The colony is homogeneous and individual phenotypes with respect to several physiological systems have been studied including neurological [38], cardiovascular [15,39–41], and endocrine [5,42]. Unfortunately, human studies are sometimes based on data from subjects with comorbidities such as diabetes and cardiovascular disease. Third, by using IHC we were able to separate measures in ZF and ZR, enabling important comparisons to better understand the impact of aging on these multiple functions of the adrenal gland. This is not possible when the whole adrenal cortex is homogenized for Western blot analysis. The major weakness of the study is that only females were available. This is often the case in nonhuman primate studies as primate centers do not maintain many males across the life course. We are have been working to assemble male baboons across the life course to rectify this limitation in the future. While these data are important in determining overall ZF and ZR control mechanisms, they do not address the ability to respond to acute environmental stimuli in these two tissues.

## ACKNOWLEDGEMENTS

We would like to thank Karen Moore for administrative support and to Dr. Edward Dick, Dr. Patrice Frost, Dr. Corinna Ross, and all of the staff of SNPRC for their support in animal care, veterinary procedures, and facility maintenance.

